# Advancing coral micropropagation for coral restoration and reef engineering

**DOI:** 10.1101/2024.06.25.600690

**Authors:** Emily Walton, Lindsey Badder, Claudia Tatiana Galindo-Martínez, David B. Berry, Martin Tresguerres, Daniel Wangpraseurt

## Abstract

In the face of escalating threats posed by human-induced climate change, urgent attention to coral reef restoration is imperative due to ongoing reef degradation. Here, we explored the potential of generating coral micropropagates as a tool to rapidly generate coral tissue for reef restoration and reef engineering. We developed a hypersalinity-induced polyp bailout protocol and a simple attachment device to support the growth of micropropagates on commonly used restoration substrates. We found that hypersalinity induction, at a rate of < 1 ppt hr^-1^, produced healthy micropropagates of the coral *Stylophora pistillata*. The highest attachment success (∼74%) was achieved in CaCO_3_ substrate devices, which outperformed PVC (∼5%) and Portland cement (∼48%). Settled micropropagates displayed rapid growth rates on both CaCO_3_ (0.037 mm^2^/day ± 0.002 SE) and PVC (0.057 mm^2^/day ± 0.008 SE) substrates, while Portland cement induced tissue degradation. Our study provides a detailed methodology for reliably generating, attaching, and growing coral micropropagates and underscores the potential of polyp bailout as a viable technique supporting coral restoration and reef engineering efforts.

## Introduction

Reef-building corals are the primary architects of tropical reef ecosystems that harbor a diversity of life forms spanning from microorganisms to large marine vertebrates (Hughes et al., 2017). Coral reefs are a hotspot for ocean biodiversity and provide crucial goods and services via fisheries, shoreline protection, nutrient cycling, and tourism activities (Eliff and Iracema., 2017; Rädecker et al. 2015). However, coral reefs are degrading at an unprecedented rate due to anthropogenic stressors such as climate change, overfishing, and pollution (Hoegh-Guldberg et al., 2017; van Hooidonk et al., 2016; Pandolfi et al., 2003).

The ongoing decline of coral reefs highlights the urgency of comprehensive conservation and restoration efforts (Suggett et al., 2023; Roger et al., 2023a) including coral gardening approaches and the development of artificial or hybrid reef structures (Burt et al., 2009; Lima et al.,2019; Rinkevich, 2021; Roger et al., 2023b). Coral gardening (Rinkevich, 2021) involves the cultivation of coral fragments under protected conditions in nurseries until they reach a certain size and maturation, and their subsequent outplanting onto existing degraded reefs (Forrester et al., 2012). Artificial and hybrid reefs (A&HR) involve the deployment of human-made structures that mimic the structure of natural reefs (Burt et al., 2009; Lima et al., 2019). There has been a growing interest in the deployment of A&HRs since the 1980s, as these structures reduce coastal erosion and enhance eco-tourism activities (Rossi and Rizzo, 2020; Higgins et al., 2022). A key challenge for creating A&HRs is to rapidly cover the man-made reef structures with beneficial benthic coral reef communities, including stony corals. Space competition of bare substrate is high on coral reefs and it has thus been of interest to rapidly cover or ‘re-skin’ bare substrates with live corals. By creating small fragments, usually between 1-4 cm^2^ in size, one can generate rapid two-dimensional spreading of coral tissue at rates several orders of magnitude higher than under standard field conditions (Forsman et al., 2015, 2017). Such micro fragments are now widely used in restoration activities (Knapp et al., 2022; Tortolero-Langarica et al. 2020; Mostrales et al. 2022). However, as corals are modular animals that are composed of repeated ‘building blocks called polyps that can bud off asexually, it is theoretically possible to seed and propagate individual polyps.

A potential option for propagating individual polyps is via polyp bailout, a natural response to stress that results in the separation of individual polyps from the coral colony (Sammarco, 1982). This process serves as an escape response to extreme stressors such as hypersalinity, extreme high and low temperatures, and acidification (Kvitt et al., 2015; Shapiro et al., 2016). Importantly, bailed out polyps retain their symbionts and can re-attach themselves to a substrate and grow into a new colony (Shapiro et al., 2016; Chuang & Mitarai, 2020). Polyp bail out has been used to create the coral-on-a-chip system, a promising microscale platform to study coral biology in the laboratory (Shapiro et al., 2016, Pang et al., 2020). While this system has been used to characterize the reattachment of bailed out polyps on a glass surface and to monitor subsequent individual polyp growth (Shapiro et al., 2016), the potential of using polyp bailout as a method for reef restoration remains largely unexplored.

Polyp bailout typically produces dozens of micropropagates from a singular coral fragment (Schweinsberg et al., 2021; Shapiro et al., 2016), an output that is much larger than traditional fragmentation methods. Thus, coral bailout holds great potential as a method to rapidly re-skin A&HR substrates or aid in the production of coral fragments for gardening applications. The aims of this study are to develop a reliable technique for rapidly generating coral micropropagates and inducing successful settlement/attachment and growth of micropropagates on commonly used restoration substrates. Specifically, we focused on the model species *Stylophora pistillata* and evaluate attachment success and tissue growth rates in response to CaCO_3_, polyvinyl chloride (PVC), and Portland cement.

## METHODS

### Corals

Colonies of *S. pistillata* were cultivated in experimental aquarium at Scripps Institution of Oceanography (UC San Diego, USA), which provides flow-through seawater at constant temperature (25 °C); an aquarium heater (EnjoyRoyal, USA) was used to further maintain temperature in each tank. Downwelling irradiance was provided at 100 μmol photons m^-2^ s^-1^ at a 10hr/14hr light-dark cycle that included moonlight simulation (4h during dark cycle) (Orbit Marine LED Current Loop). For polyp bailout experiments, we used fragments of *S. pistillata* that were about 2 cm in length.

### Induction of polyp bailout

Polyp bailout was evaluated throughout a range of hypersalinity regimes and experimental protocols. To create different salinity gradients over time, we evaluated the use of a peristaltic pump, manual adjustments in water salinity, as well as natural evaporation of water for different water volumes. Preliminary experiments with manual increases in salinity with the addition of high salinity water over a set time and the use of a peristaltic pump resulted in inconsistent salinity gradients and proved difficult to replicate. Thus, we focused on creating salinity gradients induced by natural evaporation of water. Six different volumes (25 mL, 50 mL, 75 mL, 100 mL, 150 mL, 200 mL) of filtered natural seawater (0.35 µm) (FSW) at 35 PPT were evaporated in a small glass container (8 cm diameter) to create different rates of salinity increase over time. Evaporation experiments were performed in a 25 °C room with the container placed on a magnetic stirrer at 40 rpm to create gentle water movement.

Incident irradiance was provided by a light emitting diode (LED) panel that delivered a downwelling irradiance of 100 μmol photons m^-2^ s^-1^. An air pump was connected to a small pipette to ensure that the O_2_ content of the ambient water was air saturated. Salinity was measured hourly using a refractometer (Agriculture Solutions, USA). For each salinity gradient, we exposed 3 fragments to hypersalinity stress. Each fragment was visually monitored for polyp detachment. *S. pistillata* required gentle agitation for the polyps to be released from the skeleton. This was done by utilizing a plastic pipette to gently agitate the fragment with water. The generated coral micropropagates were then transferred to a petri dish filled with 35 ppt seawater to re-acclimate to normal seawater conditions.

### Viability of coral micropropagates

The health and viability of micropropagates was assessed by examining polyp morphology using a stereoscope (Olympus SZ61, Japan) and PSII photochemistry via pulse amplitude modulated (PAM) chlorophyll (chl) *a* fluorescence imaging (Imaging-PAM). We categorized a healthy polyp morphology based on the presence of intact tentacles, a well-defined tissue structure surrounding the mouth opening, and the presence of the stomach (Schweinsberg et al., 2021). Polyps that were characterized as degraded were lacking one or more of these characteristics. Viable and healthy polyps also often demonstrated a spinning behavior, as noticed in other studies (Supplementary File S1, Shapiro et al., 2016).

For chl *a* fluorimetry, we used an imaging pulse amplitude-modulated chl *a* fluorometer (Imaging PAM, mini version; WALZ GmbH, Effeltrich, Germany) that employs a blue measuring light (460 nm). Following a dark acclimation period of 20 min, we measured the maximum quantum yield (F_V_/F_m_) of photosystem II (PSII) following a saturation pulse (Wangpraseurt et al. 2019a). We also measured relative electron transport rates (rETR) of coral micropropagates during the attachment period to artificial substrates, to evaluate the light-use efficiency (α) and maximum relative electron transport rate (rETR_max_) over time (Ralph et al. 2005). For this, rETR curves were performed spanning an irradiance regime from 0 to 783 μmol photons m^-2^ s^-1^ (0, 1, 23, 43, 81, 145, 222, 269, 321, 403, 492, 783 μmol photons m^-2^ s^-1^) with an incubation period of 30 s at each light step (Ralph et al. 2008). Five micropropagates from each substrate were measured at days 7, 14, and 21 following bail out. Data was fit to the Platt equation (Platt et al., 1980).

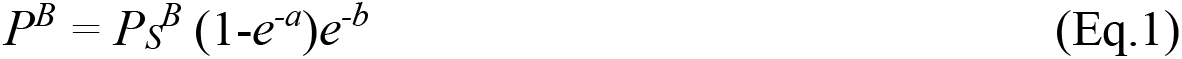

### Attachment of coral micropropagates

For our attachment tests, we used tiles made from CaCO_3_ (Ocean Wonders, USA), PVC and Portland cement (HardieBacker, USA). CaCO_3_ tiles were left at manufactured dimensions (3.2 cm x 3.2 cm) while PVC and Portland cement tiles were cut to 10 cm x 5 cm in size. Preliminary attachment experiments suggested that the creation of a small crevices improved attachment success, and we therefore fabricated crevices that were 2 mm deep and 1 mm wide using a handsaw.

To ensure sufficient gas and nutrient exchange between the micropropagates and the ambient water and to provide replication, we developed 18 laminar flow chambers (LFCs) (12 cm x 6 cm x 3 cm). These LFCs were designed using CAD software (OnShape) and 3-D printed (PRUSA MK4 3D Printer) using black PLA filament. LFCs were fitted with laminar flow dividers (6 cm x 3 cm) that were laser cut from acrylic paneling. The LFCs were then placed on top of holding tanks filled with 10 liters of FSW (35 ppt) that provided flowthrough aerated seawater at a laminar flow velocity of 1 cm s^-1^. Flow velocity was measured via tracking the movement of small particles in the water. The water temperature was set to 25 °C via a submersible aquarium heater (EnjoyRoyal, USA).

To facilitate attachment to the restoration substrates, 7 micropropagates were carefully dispersed within the crevice of each tile. Attachment experiments were performed with a total of 6 tiles distributed over 3 tanks per treatment (PVC, CaCO_3_, Portland cement), resulting in a total of 126 tested micropropagates in 18 laminar flow chambers. Experiments were performed with FSW at 25 °C, 35 ppt salinity, and an incident irradiance of 100 μmol photons m^-2^ s^-1^ (10hr/14hr light-dark cycle). To maintain salinity levels, FSW was manually added to compensate for any evaporative losses. Attachment experiments were performed only with healthy micropropagates, as defined based on morphological characteristics and Fv/Fm values > 0.3 (see above). Attachment was observed for 7 days and categorized as attached, detached or degraded (polyp lysis).

### Micropropagate growth rate

To determine the growth rate of attached micropropagates, close-up images were taken over 14 days using a stereoscope (Olympus SZ61, Japan). To minimize physical disturbance, images were taken directly in the LFCs. Lateral tissue growth was approximated using ImageJ (version 1.53, USA) by manually segmenting the planar surface area (SA, mm^2^) covered by coral tissue. The growth rate of micropropagates was analyzed as the % change in the SA of the micropropagates over a 14-day growth period.

### Optical coherence tomography imaging and substrate characterization

Optical Coherence Tomography (OCT) imaging was used to non-invasively characterize the surface structure of the artificial substrates and the attachment of coral micropropagates. OCT imaging was performed as described previously (Wangpraseurt et al., 2017, 2019b) using a Ganymede spectral domain OCT system (GAN311C1, 930 nm) with an axial resolution of 5.5 µm, an imaging depth of 2.9 mm and a lateral resolution of 8 µm (in air). Briefly, bare substrates were imaged in a black acrylic chamber filled with seawater. To calculate surface roughness, 3D OCT images were binarized in ImageJ (NIH, Bethesda, MD) and imported into Matlab (Mathworks, Natick, MA) as a tiff stack. The voxels on the surface of the substrate were identified as the binary voxel closest to the probe for each stack of voxels. Then the surface voxels were fit to a plane by simple linear regression. The difference between the surface and the best-fit plane was calculated for all points. Surface roughness is reported as the mean difference between the surface and best-fit plane for each sample.

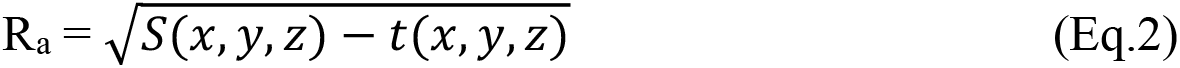

Wherein R_a_ is surface roughness, *S(x*,*y*,*z)* is the surface, and *t(x*,*yz)* is the best-fit plane.

### Statistical analysis

All statistical analyses were performed using Origin Pro (2023) and Microsoft Excel (2022). We used an alpha level of 0.05 to determine statistical significance.

## Results & Discussion

### Hypersalinity-induced micropropagates

We successfully generated viable *S. pistillata* bailed polyps using a simple evaporation-based hypersalinity gradient (Fig. 1A). These bailed polyps looked intact at salinity rates of 0.9 ppt/hr or less, while tissue disintegration occurred at faster rates (Fig. 1 A). At a salinity increase rate of 0.9 ppt/hr, > 82% of micropropagates appeared healthy as indicated by intact tentacles, stomach and mouth opening (Fig. 1A). Variable chlorophyll *a* fluorescence imaging revealed no significant difference in the maximum quantum yield (F_v_/F_m_) of photosystem II (PSII) of micropropagates compared to intact coral fragments (0.45 ± 0.04 vs. 0.47 ± 0.08; ANOVA, p = 0.208; Fig. 1 C-E). The sequence of morphological changes over the salinity gradient was reproducible and we typically observed polyp retraction when salinities of 44-46 ppt were reached, which was followed by the separation and thinning of the coenosarc at about 48-51 ppt, and total separation of polyps from the connecting tissue and skeleton at about 52-56 ppt. This process resulted in the total bailout of polyps after a period of 24-26 hours.

**Figure 1.**
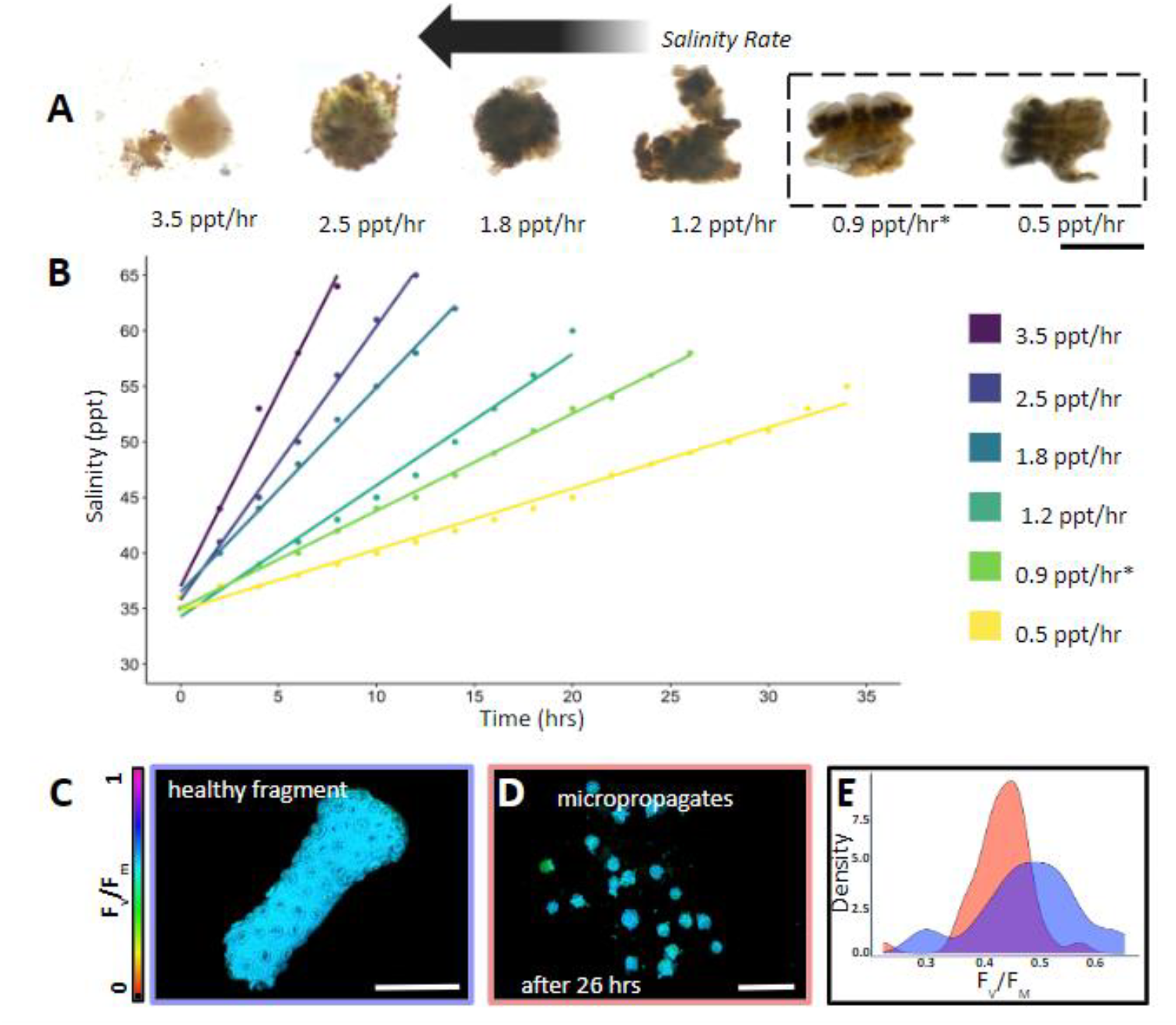
*Stylophora pistillata* micropropagate viability. **(A-B)** Generated micropropagates in response to evaporation-based hypersalinity gradients (scale bar = 1 mm). (**C-D**) Representative images of the maximum quantum yield (F_v_/F_m_) of photosystem II (PSII) of healthy coral fragments and generated coral micropropagates following 26 h of hypersalinity treatment (scale bar = 0.5 cm). (**E**) Histogram of F_v_/F_m_ of healthy *S. pistillata* polyps from intact fragments (dark blue) and generated micropropagates (red) induced via a salinity gradient of 0.9 ppt/ hr (*n*= 20 polyps).

Our study builds upon the work of Shapiro et al. (2016), which demonstrated that two other Pocilloporidae coral species (*Pocillopora damicornis* and *Seriatopora hystrix*) responded to hypersalinity stress with viable polyp bailout. However, our study developed a simple and replicable method protocol for inducing polyp bailout in *S. pistillata*. Consistent with Shapiro et al. (2016), we observed polyp bailout occurring at salinity levels 17-21 ppt above that of ambient seawater. It is noteworthy that, in contrast to previous observations in *P. damicornis*, the induction of viable micropropagates in *S. pistillata* required agitation and water circulation, presumably to alleviate hypoxic stress during darkness (Wangpraseurt et al., 2012). The similarities in polyp bailout responses across *P. damicornis, S. hystrix*, and *S. pistillata* suggest that this may be a common response to hypersalinity stress in corals, or at least within the Pocilloporidae family.

### Micropropagate attachment to coral reef restoration substrates

The use of coral micropropagates for restoration has been hampered by the inability of micropropagates to successfully attach and grow over extended periods of time (Schweinsberg et al., 2021). Here, we developed a simple attachment device that uses an aerated, laminar flow- through system equipped with micropropagate chips (Fig. 2A-D, see methods) to successfully induce attachment of *S. pistillata* bailed out polyps.

**Figure 2.**
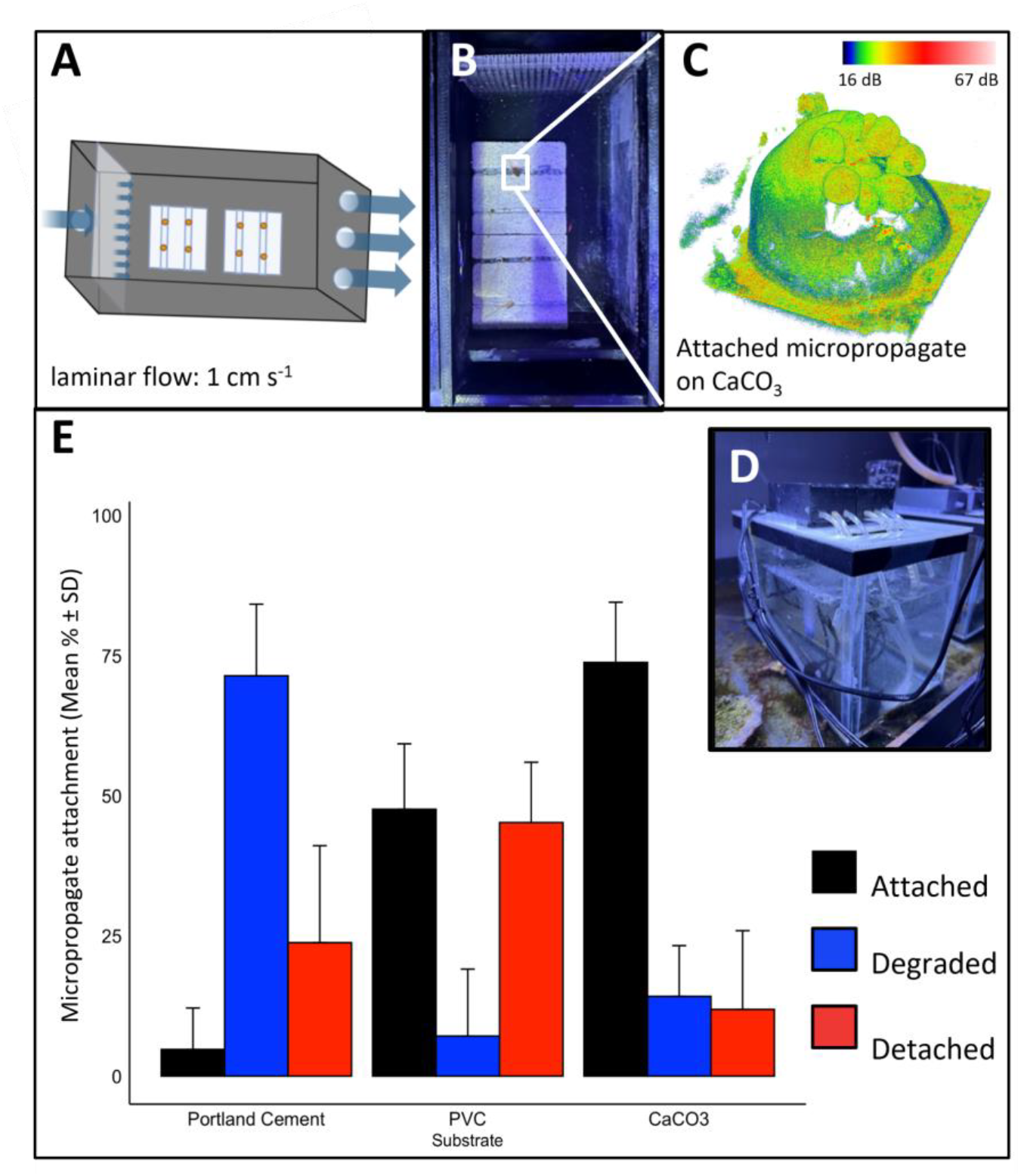
Attachment of micropropagates to restoration substrates. **(A-D) Micropropagate chip device. (A-B)** Schematic and image of the 3D printed acrylic chambers. Slow, laminar flow of aerated seawater is delivered to the micropropagates. Crevices (1 mm in width and height) facilitate tissue attachment. (**C**) Example 3D optical coherence tomography image of attached micropropagate and **(D**) image of entire recirculating system. (**E**) Micropropagate attachment success for chips made from Portland cement, PVC and CaCO_3_.

We evaluated the suitability of commonly used restoration materials, including CaCO_3_, PVC, and Portland cement as micropropagate chip substrates (Adey et al., 1975; Boström-Einarsson et al., 2020; Levenstein et al., 2022). CaCO_3_ is often used as a substrate in coral gardening efforts and is a preferred substrate as it is also the building material of coral skeletons (Levenstein et al., 2022). PVC-based substrates have been extensively used in coral nurseries and the formation of AR frameworks, due to their low cost and ease of installation. (Mallela et al., 2017). Likewise, cement serves as an inexpensive and scalable substrate that has become increasingly popular in the development of ARs (Boström-Einarsson et al., 2020; Burt et al., 2009). We found significant differences in settlement and attachment rates between CaCO_3_, PVC and Portland cement using our custom-made micropropagate attachment device (Two-way ANOVA, p = 2.39E-10 (Fig. 2A-D). The highest attachment rate was observed for CaCO_3_ (73.81 % ± 10.7 %) which was > 15-fold higher compared to Portland cement (4.76 % ± 7.38 %) and about 1.4- fold higher than for PVC (47.62 % ± 11.66 %, Fig. 2E). For the first seven days, micropropagate degradation was highest for Portland cement (71.4 % ± 12.8 %), which was nearly 5-fold greater than for CaCO_3_ (14.2 % ± 9.1 %) and 10-fold greater compared to PVC (7.1 % ± 12.0 %, Fig. 2E). Together, these results underscore the importance of material choice for optimizing attachment success of micropropagates.

We hypothesized that the observed differences in attachment success are related to surface roughness and substrate microarchitecture. We therefore used OCT, an optical analog to ultrasound (Jaffe et al., 2022), to characterize the 3D surface structure of restoration substrates with micron resolution. We found no clear relationship between microscale surface roughness (*Ra*, see methods) p = 0.287 and attachment success, (Fig. 3D, S3). *R*_*a*_ was highest for CaCO_3_ (18.5 μm ± 1.3 SE) which was 1.5-fold higher compared to Portland cement (12.2 μm ± 1.5 SE) and about 6.5-fold greater than for PVC (2.8 μm ± 0.3 SE) (Fig. 3A-D). Although it is known that larger roughness elements (e.g. mm scale crevices, Randall et al., 2021), provide protection and can enhance attachment success in a flow-driven environment, smaller roughness elements analyzed in this study (Ra < 20 μm) did not affect micropropagate performance. These results suggest that other factors have a more prominent role in modulating attachment success and micropropagate degradation within the scope of our experiment. For instance, Portland cement is known to be highly alkaline (pH ∼13, e.g. Guilbeau et al., 2003) and such enhancements in surface pH could have negative effects on coral growth.

**Figure 3.**
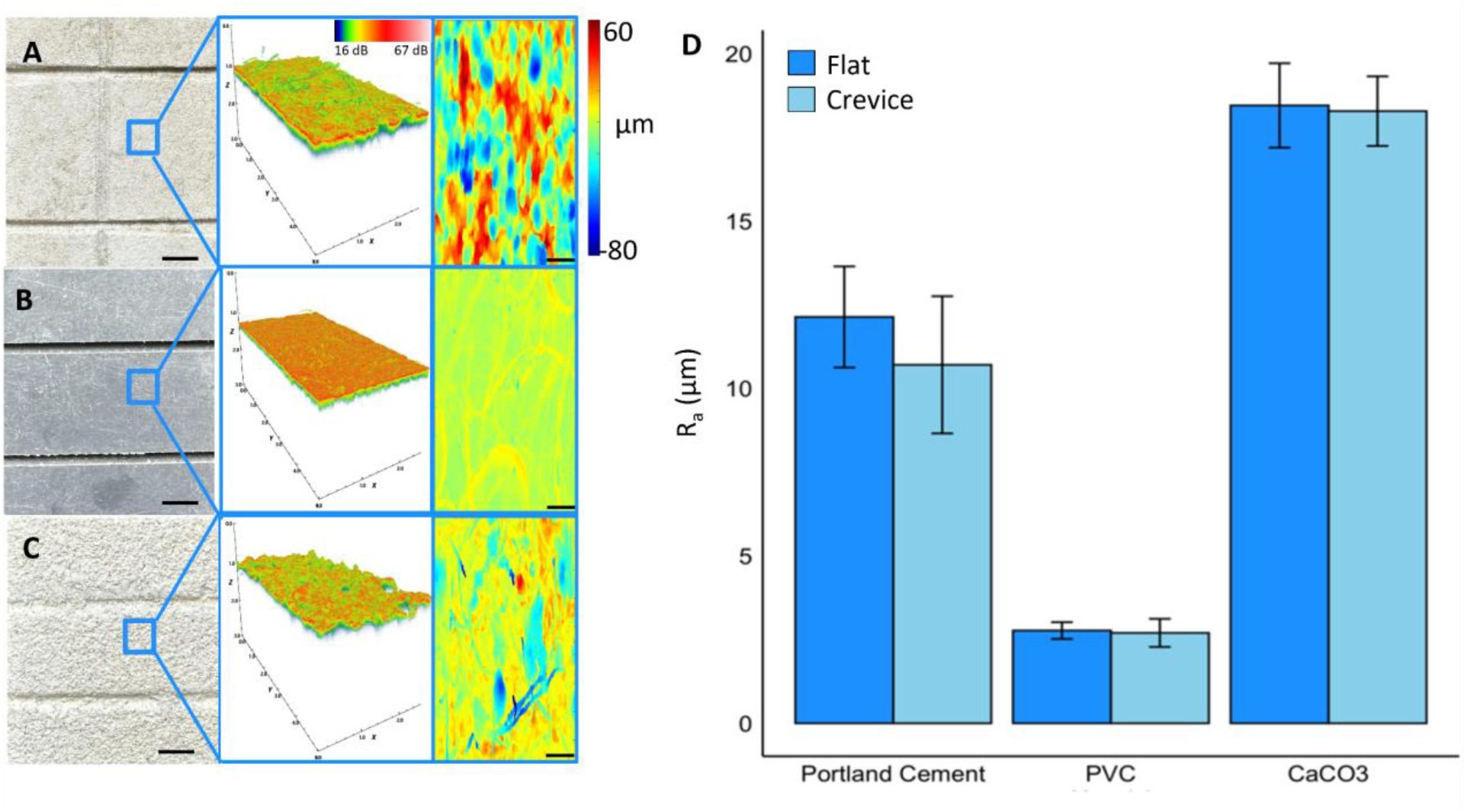
Optical coherence tomography imaging and microstructural characterization of restoration substrates. (**A**) Portland cement (**B**) PVC and (**C**) calcium carbonate substrates used in attachment experiments. Top view image (left, scale bar = 5 mm) and associated 3D OCT scan of area highlighted in blue (middle). Analyzed 2D projections of the difference in distance between the tissue height and a plane fit to the tissue surface (right, scale bar = 100 μm). (**D**) Surface roughness (*R*_*a*_) for the tested restoration substrates.

### Micropropagate tissue growth and healthiness

Using our micropropagate chip device, we successfully created growing micropropagates on PVC and CaCO_3_ chips (Fig. 4A-I). We observed the highest growth rate of micropropagates on PVC (0.057 mm^2^ ± 0.008 SE) which was 1.5-fold higher than for CaCO_3_ (0.037 mm^2^ ± 0.002 SE, Fig. 4E,H). In contrast, micropropagates growing on Portland cement experienced significant tissue degradation and did not grow laterally (Fig. 4A,B). Variable chl *a* fluorescence imaging revealed highest F_v_/F_m_ values for day 7, day 14, and day 21 for micropropagates growing on CaCO_3_ (Day 7 = 0.49 ± 0.06, Day 14 = 0.46 ± 0.09, Day 21 = 0.48 ± 0.08) which was nearly 1.5-fold higher than for PVC (Day 7 = 0.36± 0.04, Day 14 = 0.38 ± 0.03, Day 21 = 0.35 ± 0.05) and almost 2- fold greater compared to Portland cement (Day 7 = 0.28 ± 0.04, Day 14 = 0.22 ± 0.06, Day 21 = 0.20 ± 0.3) (Fig. S2). Analysis of rETR showed an increase in both the initial slope of the curve (α- light use efficiency) and the rETR_max_ over time for CaCO_3_ and PVC, eventually yielding rates similar to healthy control fragments (Fig. 4 C,F,I, Table 1). In contrast, micropropagates on Portland cement did not recover and showed continuous loss of photosynthetic potential over time (Figure 4C, S2).

**Figure 4.**
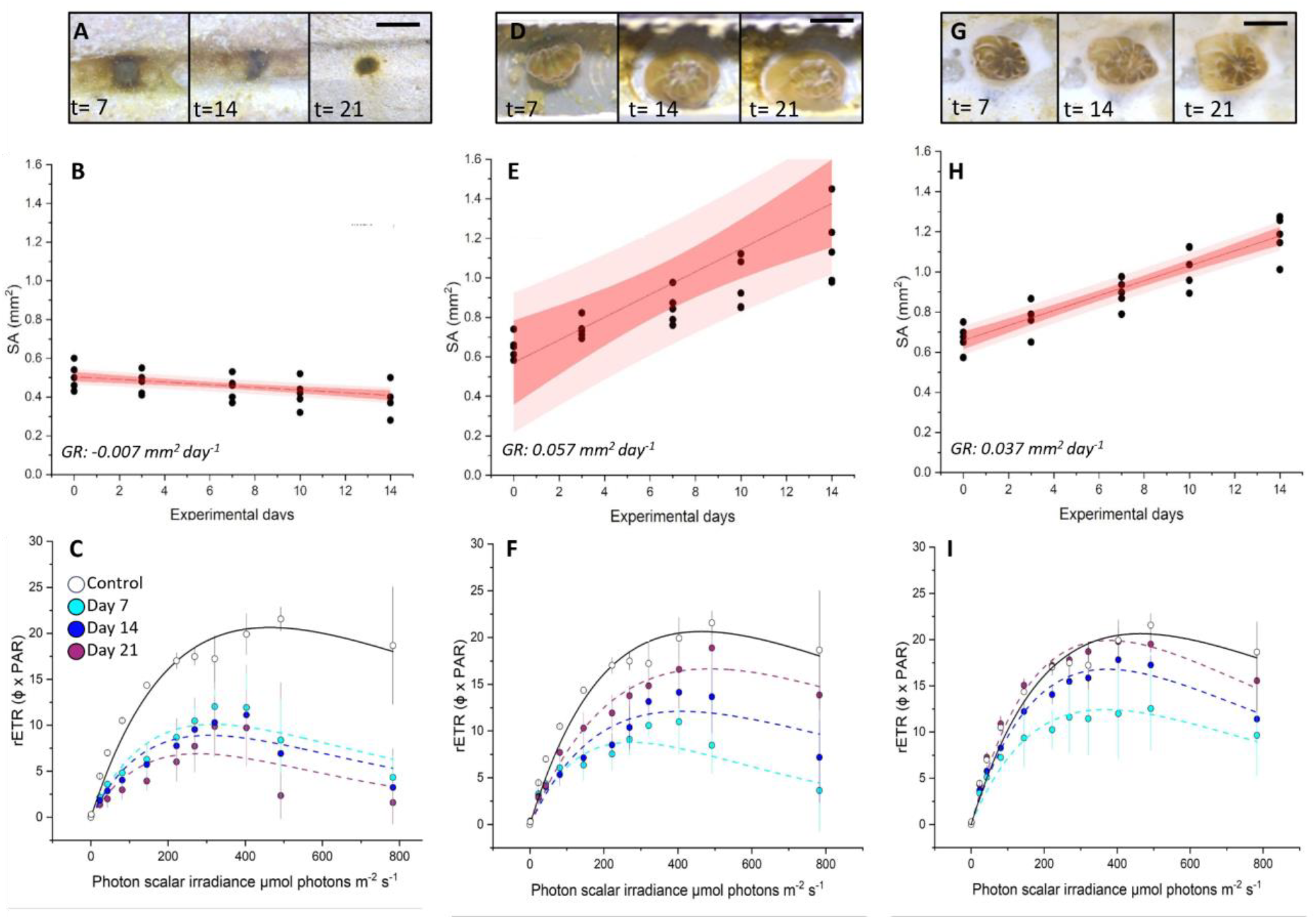
Growth rates and viability of settled coral micropropagates for (A-C) Portland cement, (D-F) PVC and (G-I) CaCO_3_ substrates. Microscope images of morphological changes at day 7, 14 and 21 (**A, D, G**). Planar tissue surface area changes (mm^-2^) per micropropagate over 14 days (**B**,**E**,**H**). Best fits (solid red line), 95% confidence intervals (red area), 95 % prediction intervals (light red area) and average growth rates (GR) are shown (*R*^*2*^= 0.93, 0.93 and 0.99 for B, E and H, respectively). (**C, F, I**) rETR of micropropagates after 7, 14 and 21 days of cultivation in micropropagate chip set-up. Control measurements (healthy fragments) were performed before bailout. On Day 21 at PAR of 492, CaCO_3_ = 19.5 ± 0.9, PVC = 18.9 ± 2.2, Portland cement = 2.3 ± 2.5. Data are means ± SE (*n=* 45). Best fits (dotted and solid lines) are based on the Platt equation (see Eq. 1).

### Applications for reef restoration

The results of this study highlight the potential of the polyp bailout approach to enhance the production of viable polyp micropropagates, underscored by the ability to generate dozens of individual polyps from a single small coral fragment (Fig. 1). We developed a simple, transferable method to generate and grow micropropagtaes from *S. pistillata* in laboratory set-ups. Using a custom-made micropropagate chip device, our results suggest that attachment and growth are substantially affected by substrate material properties, whereby CaCO_3_- or PVC-based substrates are more suitable than cement. However, the success of this technique for coral outplanting and gardening applications will strongly depend on competitive pressure from adjacent benthos. Indeed, and similar to settled coral larvae, micropropagates may be very susceptible to overgrowth by competitive macroalgae (e.g. turf algae) (e.g. Lirman et al. 2000). Therefore, the choice of substrate should also consider settlement and growth of competitive organisms (Leonard et al., 2022). A potential approach is to use micropropagates obtained by coral bail out in *ex situ* coral nurseries and leverage their rapid initial tissue growth rate, similar to current microfragment cultivation approaches (Mostrales et al. 2022). Outplanting of tissue-covered chips could then supplement the re-skinning of artificial or hybrid reefs.

In conclusion, polyp bailout offers the potential for large-scale production of viable micropropagates, representing a valuable addition to coral propagation efforts. Future research should aim to assess the performance of micropropagates produced through this technique under different environmental conditions and how micropropagates compare to other fragmentation techniques under *in situ* conditions. This will contribute to a more comprehensive understanding of the advantages and limitations of polyp bailout in the broader landscape of coral restoration strategies. Additionally, due to their similar properties to settled coral larvae, micropropagates can also be used as a suitable model system for studying mechanisms that control post-settlement mortality (e.g. competition, predation, detachment).

## Acknowledgements

This study was funded by grants from the US National Science Foundation (NSF-BSF IOS award # 2149925 to DW and 2149926 to MT, award NSF DBI award # 2316391 to DW) as well as the Defense Advanced Research Projects Agency under the Reefense Program, BAA HR001121S0012 to D.W. The views, opinions, and/or findings expressed are those of the authors and should not be interpreted as representing the official views or policies of the Department of Defense or the US government.

## Author Contributions

Designed the project: EW, DW, MT. Performed experiments: EW, LB, CTG. Analyzed and interpreted data: EW, LB, CTG, DBB, MT, DW. Supervised the study: DW, MT. wrote the paper: EW and DW with editorial contributions from all authors.

## Conflict of Interest

The authors declare no conflict of interest.

## Notes

### Competing Interest Statement

The authors have declared no competing interest.

