## Supplementary Information for "Advancing coral micropropagation for coral restoration and reef engineering"

Walton et al.

**Supplementaty File S1:** Video of spinning of coral micropropagate

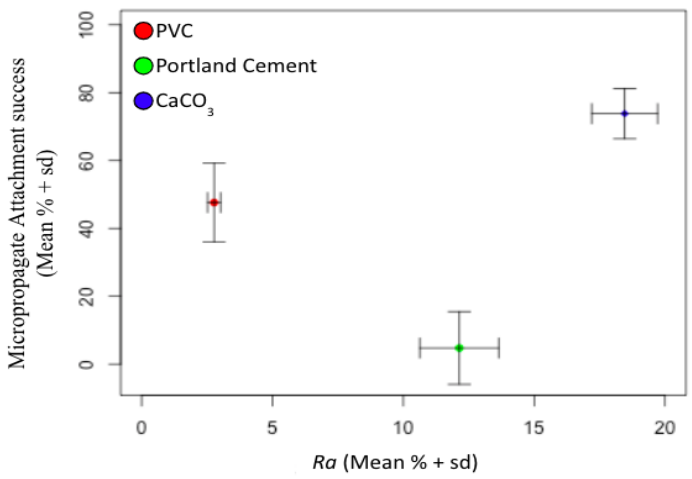


**Fig. S1. Microcropagate attachment success vs. surface roughness ($R_a$).** ($CaCO_3$ (Average Roughness = 18.4 ± 1.0 , Attachment success = 73.8 % ± 10.8 %) Portland cement (Average Roughness = 11.4 ± 1.8 , Attachment success = 4.8 % ± 7.4 %) PVC (Average Roughness = 2.7 ± 0.3 , Attachment success = 47.6 % ± 11.7 %), *n* = 36)

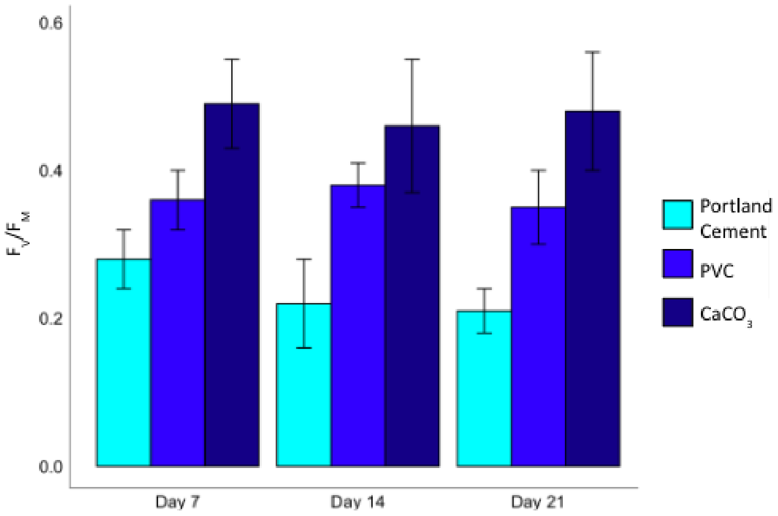


**Fig. S2. $F_v/F_m$ of micropropagates grown on Portland cement, PVC and $CaCO_3$** ($CaCO_3$ (Day 7 = 0.49 ± 0.06 , Day 14 = 0.46 ± 0.09, Day 21 = 0.48 ± 0.08) PVC (Day 7 = 0.36± 0.04 , Day 14 = 0.38 ± 0.03, Day 21 = 0.35 ± 0.05) Portland cement (Day 7 = 0.28 ± 0.04 , Day 14 = 0.22 ± 0.06 , Day 21 = 0.20 ± 0.3), *n*=45)
